# Experimental Emergence of Conventions in Human Dyads

**DOI:** 10.64898/2026.01.10.698799

**Authors:** Oviya Mohan, Dora Biro

**Affiliations:** Department of Brain and Cognitive Sciences, University of Rochester, New York

**Author notes:** Corresponding author: Oviya Mohan.

**Keywords:** coordination, convention, social cognition, strategic behavior, problem-solving

## Abstract

Conventions can be defined as arbitrary and self-sustaining practices that emerge in a population and facilitate solving coordination problems. A recent study traced the formation of simple conventions in captive baboons in a touch-screen-based color-matching ‘game’. We replicated this task with human pairs under different conditions (with/without visual access to the partner’s screen; with/without prior information on the task structure) to assess their effects on the formation and stability of conventions. We found that more information delayed the formation of conventions (arbitrary rankings of colors that determined choices in any given color-pairing). Analysis of self-reported strategies did not reveal a clear effect of condition on levels of elicited strategic behavior. Interestingly, pairs maintained their conventions even when given visual access to their partner’s screen, despite the availability of an alternative, potentially simpler, cognitive strategy. In a follow-up variant of the task that paired up experienced subjects with a naïve partner, conventions emerged faster but did not replicate the convention previously formed by the experienced player, demonstrating the transmission of “know-how” but not “know-what” information. We discuss the implications of our results for understanding the cognitive mechanisms necessary to support the formation, maintenance, and transmission of conventions.

## 1. Introduction

Coordination problems occur widely in nature and across many domains of behavior. For example, in reproduction synchronizing courtship behaviors can influence mating success [1,2], in biparental care effective timing of the handover of duties between parents can increase offspring survival [3,4], in animal movement executing a specific collective escape response can improve a group’s chances of evading a predator [5], and in foraging the success of group hunts increases if the hunters are well-coordinated with each other [6].

Many coordination problems take the form of repeated multi-agent interactions where the same problem is encountered by varying subsets of a population (sometimes with the same members and sometimes with others) and are characterized by having more than one possible solution, which may be equally good. These repeated interactions can allow the population to converge on a specific solution if the individuals involved selectively and consistently adopt one of the possible alternatives. Following the seminal work of Lewis [7] we refer to such solutions as “conventions”: arbitrary and self-sustaining practices that emerge in a population and facilitate efficient solving of a specific coordination problem (see also [8] for a recent review). Conventions can, in turn, vary across populations since there may be multiple stable solutions to the coordination problem rather than a single “correct” or globally optimal solution – instead, which solution a population settles on may be determined by stochastic historical contingencies. For example, greeting behavior in humans can consist of shaking hands or bowing (where both parties are expected to perform the same action), and which manner a given population uses is a function of that population’s own unique history of inter-individual interactions. Similarly, driving conventions dictate whether motorists drive on the left or the right side of the road on a country-by-country basis, with either solution facilitating efficient and safe traffic flow as long as it is consistently adopted by the population at large.

Conventions are characterized by three key properties [9]. First, they are *efficient*: once established in a population, each encounter with the same problem can be solved quickly, even if coordination partners vary (but only as long as these partners share the same convention). Second, conventions are *arbitrary*: the solution can take multiple forms, where one is not necessarily superior to another, and hence different populations may converge on different solutions. Third, conventions are *self-sustaining*: no individual benefits from deviating from the established convention on their own and attempts to shift the convention at the population level can have disastrous consequences (the massive planning operation that accompanied Sweden’s overnight switch in 1967 from driving on the left to driving on the right represents a good example).

The emergence of conventions has been explored across several disciplines including linguistics, behavioral economics, game theory, computational social sciences, philosophy and ethology yielding a variety of insights into the mechanism of when and how conventions emerge [7,10–15]. Some approaches have focused on how external factors such as problem structure, solution space, or the network configuration of interacting individuals (e.g., [16–18]) impact the emergence of conventions. Other approaches have focused on exploring what higher-level cognitive processes scaffold the emergence of conventions at the population level (e.g., [7,12,19]), while yet others have explored how conventions scale up from small-scale interactions to the population (e.g., [20,21]). Recently, cognitively inspired computational models have aimed to understand the role of individual-level processes such as social learning (e.g., [22]) and reinforcement learning (e.g., [23]) in the emergence of conventions. While computational models with artificial agents have great potential for scaling up to investigations at the level of large population and transmission over multiple generations, empirical human data is necessary to obtain better informed and sufficiently complex models so as not to fall into the trap of being too simple (e.g., Rogers’ paradox, [24]). To build better models, there are still gaps in our framework for understanding conventions, specifically the cognitive mechanisms at play at multiple levels, that require empirical data [10,25,26].

As outlined in recent reviews [8,26], conventions are theorized to emerge via multiple mechanisms operating at three different levels. At the individual level, basic cognitive abilities like learning and memory as well as more complex components of social cognition like imitation and reasoning about others are operational. At the group level (i.e., some subset of the population, with the smallest unit consisting of a dyad of two individuals), coordination may be based on joint goals. At the population level, demographic factors such as the rate of turnover in the population or the frequency of interactions with informed vs uninformed members of the population, or the population’s network structure, may affect the likelihood of the emergence of a population-wide convention. Wu and colleagues [26] propose that group-level processes are crucial to moving from individual level processes to population level solutions like conventions. They postulate that this occurs via two feedback loops, one between individual learning and group interactions (social learning facilitates coordination and vice versa) and the second between group interactions and large-scale societies (successful coordination facilitates shared knowledge and vice versa).

However, open questions remain about how group-level processes requiring coordination are facilitated when no societal-level shared knowledge (conventions) exist yet, potentially for a novel problem not yet encountered by the population. In these cases, we also do not know what/how individual level processes are utilized at the dyadic level and how dyadic interactions scale up to population level shared knowledge. Varying the conditions under which the same problem is presented to dyads allows manipulation of cognitive strategies that are available to the individuals as they complete a novel coordination task (e.g., [11,27]) and provides a way to gain insight into the underlying cognitive processes. For example, how a simpler mechanism like copying versus a more complex one involving setting expectations about one’s partner’s actions and intentions affect the emergence of a solution can be explored by manipulating visual access to the partner (copying is possible only when one can see the partner’s actions). Similarly, manipulating the amount of information the subjects have about the payoff allows us to modulate the desirability of achieving a joint goal for the pair. Solutions are likely to emerge quickly and efficiently when the subjects know they are required to coordinate, but how efficiently social coordination is achieved when these joint goals are not as concrete or transparent and have to be learned on the fly remains unclear.

To address these questions, we used a simple coordination paradigm involving a color matching task. This paradigm was adapted from a recent study [9] showing the formation of simple conventions in a non-human primate, the Guinea baboon (*Papio papio*). Pairs of captive baboons were repeatedly presented with two colored squares on a computer screen, chosen randomly from a total of seven different colors. Subjects played in ever-changing pairs, and they were rewarded if both members chose the same color in any given trial. Over time, the baboons formed a convention consisting of a color preference hierarchy which allowed them to solve the task at above-chance levels. Importantly, this hierarchy was shared at the population level even though subjects only ever interacted in pairs. Furthermore, while subjects initially performed this task in a setting where they could see each other’s screens, they were also able to maintain their convention when an opaque partition was later introduced, and to form conventions when the experiment was repeated with a new set of stimuli and no visual access to the partner’s screen at any point. While Formaux and colleagues’ [9] study provides a valuable framework in which to study the emergence of conventions and was able to address Wu and colleagues’ [26] second feedback loop (between dyadic processes and transmission to the population level), open questions remain about the first feedback loop (between the individual and the dyad) as mechanisms at play at the individual level are difficult to disentangle (for example, the extent to which the subjects viewed the task as a coordination task rather than an individual task where they were sampling reward contingencies) and in turn how this contributes to the second feedback loop. We therefore implemented this task with human dyads and under a variety of conditions to address how (i) varying participants’ knowledge of the nature of the coordination problem (*informed* vs *not informed*), (ii) varying participants’ visual access to their partner’s actions (*transparent* vs *opaque*), and (iii) introducing asymmetry in participants’ prior experience with the task (*experienced* vs *naïve*) affected the emergence of conventions. We expected to see conventions (efficient, arbitrary, and stable solutions) emerge more often and faster when individuals had the maximum amount of information, i.e., when they knew the task’s payoff structure, had access to their partner’s actions, and/or had prior experience with the task. By varying the order in which dyads were presented with the different conditions under (ii), and by examining how conventions that emerged in a given dyad were transmitted to naïve subjects under (iii), we could also characterize the maintenance of these conventions both within dyads as a function of visual access, and between dyads from different “generations”, respectively. In addition, we postulated that the different conditions would evoke varying levels of strategic behavior based on the cognitive complexity they present to the subjects; we used participants’ self-reported strategies to explore whether this was indeed the case. Our findings will help fill gaps in our theoretical knowledge of how different mechanisms may interact to give rise to solutions to novel coordination problems in dyads and how these solutions may transmit to and diffuse through larger populations.

## 2. Methods

### 2.1. Subjects and Experimental Procedures

101 human subjects – recruited from among undergraduate and graduate students at the University of Rochester (mean±SD age = 20.59±2.88 years; 15 males and 86 females) – participated in the study. The experiment consisted of two phases: in Phase I we examined the formation of new conventions in pairs of initially naïve subjects under a variety of conditions, while in Phase II we explored the transmission of a previously established convention from knowledgeable to naïve subjects. Written consent was obtained from all subjects, and the study was approved the by the Institutional Review Board of University of Rochester (Study ID: STUDY00007197). The recruitment period for the study was from November 11, 2022 to April 30, 2023.

### 2.2. Phase I

Eighty subjects completed a simple color-matching task adapted from Formaux and colleagues [9]. All subjects were initially naïve to the task and were assigned to pairs randomly, yielding a total of 40 pairs. Partners in a given pair were seated in front of 21.5-inch computer screens next to each other, at a distance of approx. 1 m. Each pair performed two sessions of 294 trials each, and, for the first session (Session 1), was randomly assigned to one of four conditions distinguished by (i) whether or not the subjects had visual access to each other’s screens (*opaque* vs *transparent*), and (ii) whether or not the pair was given specific information about what counted as a “correct” response in a trial (*instruction* vs *no instruction*). The four experimental conditions were constructed as different pairwise combinations of these settings, as described further below.

In the *transparent* (T) setting, subjects were able to see their partner’s screen, including the partner’s cursor and its movements, but a partition (76cm x 51cm black poster board held in place at 81cm above the floor by a metal frame) was placed between them such that they could not see their partner to minimize any non-verbal communication between the subjects (Fig 1A). In the *opaque* (O) setting, the same partition was placed such that the subjects could not see their partner or their partner’s screen (Fig 1B). In the *instruction* (I) setting, subjects were told in advance that they would score correctly on a trial (receiving 10 points) if they and their partner chose the same color; those in the *no-instruction* (NI) setting were only told that whether or not a trial was scored as correct depended on both their and their partner’s choice. The four conditions tested in Session 1 were therefore I-T, I-O, NI-T, and NI-O. For Session 2, performed immediately after Session 1, each pair’s visual access was reversed but no further instructions were provided. This meant that I-T and NI-T pairs became I-O and NI-O pairs, respectively, while I-O and NI-O pairs became I-T and NI-T, respectively. Subjects were instructed not to communicate with each other in any way throughout both sessions, and to try to maximize their cumulative score.

**Figure 1.**
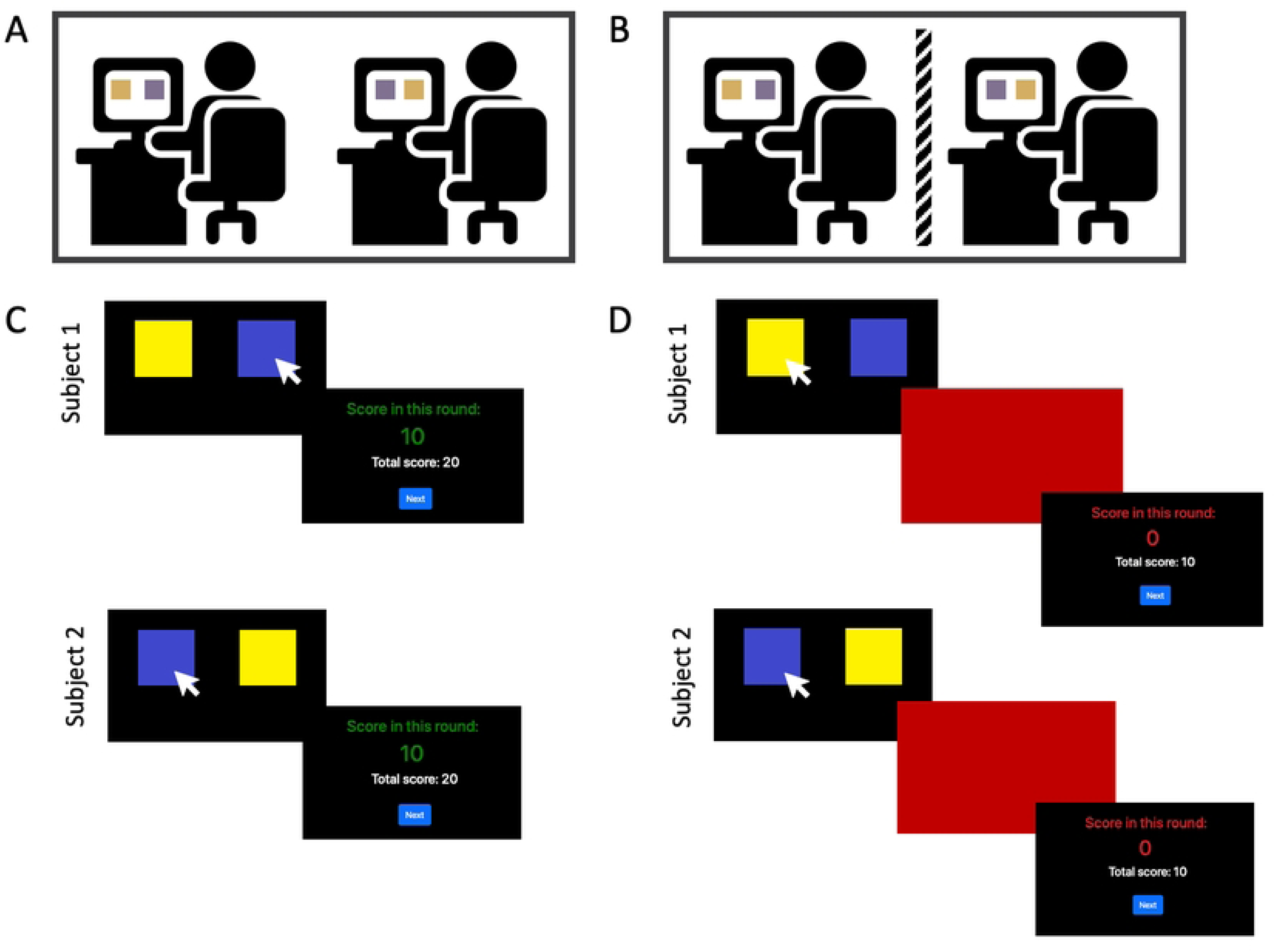
Task set-up and trial progression. Subjects and apparatus in the (A) Transparent condition where subjects can see each other’s screen but not each other and (B) Opaque condition where subjects cannot see each other’s screen or each other (icon modified from Flaticon.com). Trial progression showing both subject’s screen for a (C) correct trial where both subjects choose the same color and see the results screen showing a reward of 10 points for that trial (round) and (D) incorrect trial where the subjects choose different colors, see a brief red flash on the screen followed by results screen showing 0 points obtained for the trial (round). In both cases, total score is tracked and shown on the results screen. If one player chooses first, a waiting screen is displayed until the other player makes a choice.

The task used seven different colors (identical to those used by Formaux and colleagues [9]), offering a total of 21 different pairwise combinations. In each session, these sets of 21 combinations were presented repeatedly over 14 consecutive blocks (where the order of pairings within each 21-trial block was randomized) yielding a total of 294 trials per session. This number of trials was determined to be adequate to observe the emergence of conventions (arbitrary rankings of colors that determined choices in any given color-pairing) through pilot studies. Each session lasted ∼20-30 minutes. Subjects were prompted to take a break every 98 trials, resulting in two breaks per session.

At the start of Session 1 instructions were presented on the screen and the task began once both subjects indicated that they were ready by clicking on a start button (see online repository for exact instructions). In each trial, two colored squares (200x200px) were presented against a black background on the left and right side of the participants’ screens. The left-right locations of the colors were randomized such that in ∼50% of the trials the two colors appeared in the same location for the two subjects and in the remaining ∼50% of the trials they appeared in the opposite locations). The subjects had to make a choice by mouse-clicking on one of the squares. Either subject could respond first and whoever responded first was shown a screen indicating that the other subject had not yet responded. Once both subjects had responded, an identical feedback screen was displayed on both screens informing subjects either (i) that they were correct and would be rewarded with 10 points or (ii) that they were incorrect and would receive 0 points (see Fig 1C and 1D, respectively). Their total score up to that point was also displayed. The next trial then commenced once both subjects indicated their readiness by clicking the “Next” button.

Once a pair had completed Session 1, they were allowed to take a 5-min break, and once both subjects were ready the second session began. The same instructions page presented at the beginning of Session 1 was shown as a reminder with no new information. Crucially, the partition was moved during the break so as to switch between the opaque and transparent settings.

At the end of each session, subjects were asked to report the strategies they used to solve the task in a free-response questionnaire. For both sessions, across all four conditions, the partition was used to cover the partner’s screen from view while typing the response to ensure subjects did not see their partner’s description of their strategy.

### 2.3. Phase II

In Phase II, all subjects from pairs that used a convention in Phase I (across any of the four conditions) were invited back for a second visit. 21 subjects were able to return (7 from I-T, 6 from I-O, 3 from NI-T, and 5 from NI-O), each of whom was designated an “experienced subject” and was randomly paired with a “naïve subject” (i.e. one who had not done the task previously). Throughout Phase II, naïve subjects were not aware that they were playing with an experienced subject and vice versa.

At the beginning of Phase II, each experienced subject was asked to recall the color hierarchy they had used in their first visit to the best of their ability while their assigned naïve partner waited outside the room. The recall task consisted of the subject having to arrange the seven colors (displayed as colored squares identical to those used during Phase I) from highest to lowest ranked by dragging and dropping them into one of seven empty boxes on screen. Instructions were shown on the screen and no other verbal instructions were provided, except to answer any questions the subject may have had (see online repository for code used to create the recall task).

All pairs in Phase II then completed one session of the task in the opaque condition (O) and were given explicit instruction (I) about the task (identical I-O condition in Phase I Session 1). The duration of the task was equal to the two sessions in Phase I (total of 588 trials) completed as a single session. Subjects were prompted to take a break every 98 sessions, resulting in a total of 5 breaks. The entire session took ∼40-60 minutes. The subjects were only asked to report their strategies once at the end, after completing all 588 trials. Subjects were not allowed to communicate with each other in any way.

Throughout both Phases of the study, we used IPS (in-plane switching) monitors to ensure that the colored square stimuli appeared sufficiently similar irrespective of viewing angle (relevant only to the transparent setting where subjects were able to view each other’s screens). The main color-matching task was designed with oTree, an open source Python-based platform for web-based interactive tasks, including multiplayer games [28], and was hosted on Heroku, a cloud application platform. An experimenter (O.M.) was present in the testing room throughout all experimental sessions.

The analyses of the data concerning the calculations of the Elo-ratings, emergence, and stability of conventions were devised using pilot data from eight pairs (two assigned to each of the four conditions). All analysis code with respect these calculations was written and finalized before the collection of the data reported here, which does not include the pilot pairs. The hypothetical examples used to create the template for strategy coding were also based on strategies reported in the pilot study.

### 2.4. Analysis

#### 2.4.1. Phase I Session 1

##### Overall performance

For each pair in each session, we recorded the total number of correct responses scored, as well as the pair’s mean response times across all trials. Response times in each trial were measured from the time the colored squares appeared on the screen until both subjects had chosen a color. We also classified each pair as having either developed or not developed a convention during a given session based on whether or not the two subjects of the pair showed evidence of matching color hierarchies in their responses (see “Color hierarchies” and “Emergence of conventions” sections below), and tested whether the proportion of pairs with conventions differed across conditions using a Fisher’s exact test (fisher.test() from the R package “stats”). Accuracies and mean response times were compared between convention-forming and non-convention-forming pairs using two-tailed t-tests.

##### Color hierarchies

For each individual subject, we analyzed their pattern of responses by tracing the evolution of their preferences for each of the seven colors over the course of both sessions. In the opaque conditions (I-O and NI-O), there was only one way to solve the task with accuracies above chance level: the two subjects needed to establish a *shared* hierarchy of color preferences. This would ensure that when presented with any combination of two colors, both subjects would be able to choose the same one. In the transparent conditions (I-T and NI-T), this strategy was one of two possible solutions; the other involved the trial-by-trial use of social information (one player choosing a color and the other player copying it). This second strategy did not require a consistent color hierarchy.

To measure the emergence of color hierarchies, we calculated and tracked Elo ratings for each color trial-by-trial, separately for each subject following Formaux and colleagues [9]. Elo ratings were first developed to rank chess players [29] but more recently have also been used to construct social dominance hierarchies in behavioral ecology (e.g., [30]). The Elo-rating method continuously updates interactants’ ratings based on the outcome of each observed interaction between them (e.g. win/lose). We applied the Elo-ratings R package (R version 3.4.1, [31]; [30]) to the seven colors, separately for each subject, treating each color as an “individual”. At the start of the session, all the colors were assigned a default Elo-rating of 1000. Then, at the end of each trial the color from the current pairing that was chosen by the subject (i.e., the “winner” of the interaction between the two colors) received a boost to its Elo-rating while the Elo-rating of the color not chosen by the subject (the “loser”) decreased by the same amount. This continuous updating across the trials allowed us to visualize the evolution of color preferences for each individual subject over the course of both sessions. Figure 2A shows an example of a subject that developed a clear hierarchy of color preferences, Figure 2B shows a subject with a partial hierarchy where there was a preference for darker over lighter colors but the relative rankings of some of the colors fluctuated, and Figure 2C shows a subject who did not seem to have developed a hierarchy of color preferences.

**Figure 2.**
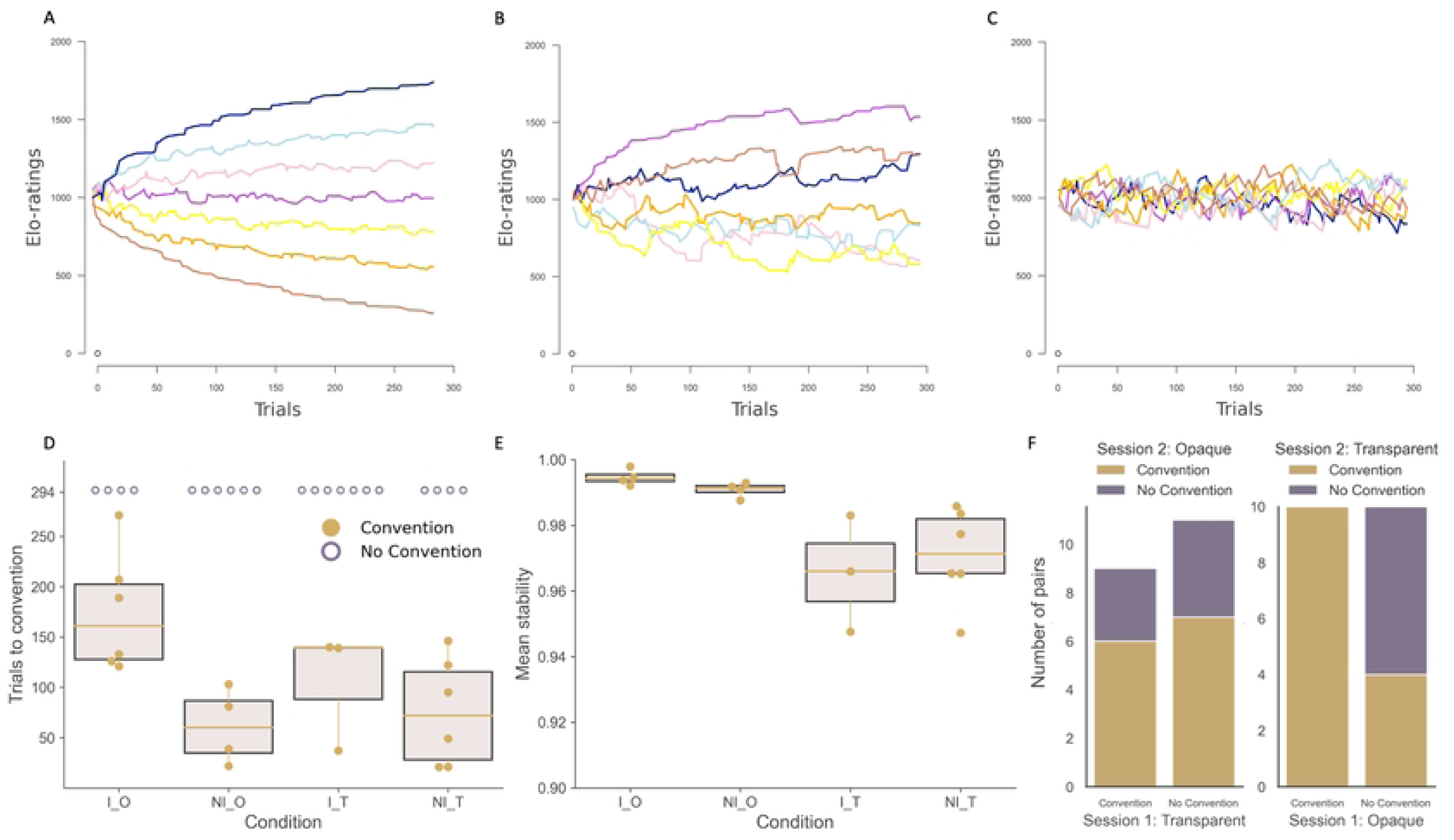
Phase I Results: Emergence, Stability and Maintenance of Conventions. Examples of the evolution of Elo-ratings for the seven colors used in the experiment (three different subjects; Session 1 only). (**A)** shows a subject with a clearly discernible color hierarchy; (B) shows a subject with a partial hierarchy (only some colors’ ranks in the hierarchy are well separated); and (C) shows a subject who did not develop a color hierarchy. Cumulative results across pairs showing (D) trials taken to reach a convention in the pairs that had a convention (solid orange dots). Hollow purple dots indicate pairs that did not converge on a convention and were not included in the calculation of the means for each condition and (E) mean stability of the conventions that emerged across the four conditions. For D and E, upper and lower bounds of box represent upper and lower quartile, respectively, center of the box represents the median and the whiskers extend to maximum and minimum (outliers are calculated ± 1.5 * inter quartile range past the high and low quartiles). Number of pairs who maintained, lost or newly developed a convention as they transitioned from (F; left) transparent session 1 to opaque session 2 and (F; right) opaque session 1 to transparent session 2.

The task allows the pairs to converge on any hierarchy of the seven colors (out of a total of 7! = 5040 different possible hierarchies). To test whether there was any universality in preferences, the average rank of each color across the hierarchies used in pairs with conventions was calculated and Kendall’s W was used to measure agreement of color order across the pairs.

##### Emergence of conventions

Qualitatively, we defined a convention as a situation in which the color hierarchies of the two subjects in a pair matched. To establish whether and when this point was reached in a given session, we defined three different measures of accuracy: *individual accuracy*, *pair accuracy*, and *combined accuracy*. *Individual accuracy* was a measure of how consistent an individual subject’s choices were with their own hierarchy of color preferences developed up to that point. It was encoded as a binary variable at each trial: 1 if the subject chose the color with the higher current Elo-rating and 0 if the subject chose the other. *Pair accuracy*, calculated for the pair at each trial, served as a measure of whether the choices of the two subjects in the pair matched (and hence whether they had solved the trial correctly). It was also encoded as a binary variable: 1 if both subjects chose the same color and 0 if they chose different colors. *Combined accuracy* combined both of these measures using a logical AND operator, i.e. combined accuracy was 1 for a trial only if both subjects had an individual accuracy of 1 and the pair accuracy was 1. This ensured that a combined accuracy of 1 could only be achieved if a color hierarchy emerged that was shared by the two subjects.

At the trial level, a combined accuracy of 1 still did not indicate a “convention” since it was possible to obtain this by chance. We used a rolling window (of size 21 trials) to track mean combined accuracy and defined a criterion above which it had to be maintained in order for the pair to qualify as having a convention. We chose a threshold of 80% so as to not penalize the pair for momentary lapses in attention and memory as well as minor confusions between color rankings. The pair had to maintain performance above this threshold for at least 21 consecutive overlapping windows (i.e., a total of 42 trials, which ensured that all unique pairings of colors were presented at least once). The first trial of these 21 windows was defined as the point of emergence of a convention for the pair. Supplementary Figure S1 shows complete data for two example pairs with panel C and G demonstrating combined accuracy as a function of trial number for a pair that reached a convention at trial 133 and a pair that did not develop a convention, respectively.

For the pairs where a convention was observed to emerge, we tested if there was a significant effect of condition on the time taken for the convention to emerge using a multiple linear regression model with instruction type (I vs. NI) and visual access type (O vs. T) as two independent variables and trials-to-convention as the dependent variable.

Assumptions about the data required for running a multiple linear regression including normal distribution of residuals and homoscedasticity were confirmed through visual inspection of the histogram of residuals, while the lack of multicollinearity of the independent variables was ensured by their orthogonal nature. If the interaction between the two independent variables was not statistically significant then the regression was rerun without the interaction term. As a follow-up if the linear model was significant, two-tailed independent t-tests were performed to determine the direction of the effects observed (lm() and t.test() from R package “stats” were used here and in following regressions and follow-up t-tests). Similar methods were used for all following linear models. All tests for statistical significance were performed with alpha set to 0.05.

##### Stability of conventions

The metric pinpointing the emergence of a convention ensures that both subjects share a color hierarchy (to a certain degree) and are able to successfully complete the task (choose the same color) for a minimum number of consecutive trials. However, this metric does not consider the stability of the convention. For example, a pair of subjects could successfully maintain their convention for the required number of trials but then experience a drop in performance if one or both subjects deviate from the established color hierarchy. Therefore, the strength or stability of the convention also needs to be evaluated. We used a measure of stability based on “crossovers” in Elo-ratings, i.e., the frequency of changes in color rankings for each individual once a convention emerges. We used a version of the stability calculations, from the Elo-ratings package [30], that did not implement weightings (weight is relevant for animal dominance hierarchies where switches in rank between two high-ranking individuals are considered to disrupt stability more than switches between two low-ranking individuals). The formula for calculating color hierarchy stability, S, adapted to our task is shown in Equation 1, where n is the trial where a convention first emerged for a pair and *X_ij_* is the Elo-ranking of color *j* in trial *i*. Values for S range between 0 and 1, with 0 being the least stable and 1 indicating no changes in rank observed. Stability was calculated for 100 trials after a convention was reached to ensure the same number of trials were included in the formula across all the pairs. Stability was calculated for each subject and averaged across the two subjects in a pair to obtain a mean stability for the pair.

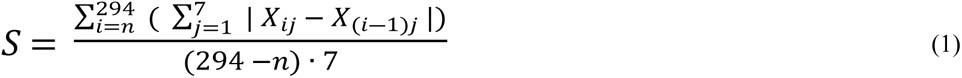

We tested whether the stability of the conventions varied as a function of condition using a multiple linear regression model with instruction type (I vs. NI) and visual access type (O vs. T) as two independent variables and the arcsine-transformed mean stability of the pair as the dependent variable.

##### Self-reported strategies

In strategic games, it is hypothesized that players assign levels of strategic behavior to their partners, employing varying levels of cognitive complexity [32]. We analyzed the subjects’ self-reported strategies by assigning them to one of three levels. Level 0 corresponded to subjects who did not indicate that they took their partner’s actions into account. Level 1 players assumed their partner was playing at level 0 and played accordingly. Level 2 players assumed their partner was at least level 1, i.e. that their partner also had a strategy that took their actions into account (Supplementary Table 1 shows hypothetical examples used by coders as guidelines and examples of subject reported strategies). The strategies were rated by two coders, one who was aware of the goals of the study and one who was not. There was substantial agreement between the two coders (Cohen’s Kappa, κ = 0.74). Ratings where there was mismatch (12 subjects) were assigned a final rating via discussion between the two coders.

For each pair, we calculated the minimum level of strategy used by the two subjects as well as the maximum level. A multiple linear regression model with instruction type (I vs. NI) and visual access type (O vs. T) as two independent variables and strategy (either minimum or maximum strategy) as the dependent variable was used to test whether the different conditions evoked the need for less or more complex cognitive strategies. These analyses were conducted only with Phase I Session 1 data for 39 out of the 40 pairs, one pair’s strategy were only partially saved due to a technical error and thus not used for this analysis. For pairs with a convention, a multiple linear regression model with instruction type (I vs. NI) and visual access type (O vs. T) as well as minimum or maximum strategy as independent variables and trials-to-convention as the dependent variable was used to test if level of strategic behavior had an effect on how quickly conventions emerged. These models were compared to the model described in the “Emergence of conventions” section as a null model (without strategies) for 18 out of the 19 pairs with conventions (as one pair’s data was lost due to a technical error).

#### 2.4.2. Phase I Session 2

##### Maintenance of conventions across conditions

Data from Session 2 were used to explore whether conventions that formed in Session 1 were maintained across a change in visual access. The same metrics as for Session 1 were calculated, with one key difference: the Elo-ratings of the colors at the beginning of Session 2 were set to the final Elo-ratings at the end of Session 1 to ensure a fair assessment of convention emergence and/or maintenance (starting at 1000 again would have invalidated the hierarchy/knowledge that the subjects had built up in Session 1).

Pairs in T and O conditions in Session 1 (I and NI were collapsed since by the end of Session 2, every pair had had a chance to play in the T setting and thus could be assumed to have understood the task reward/payoff structure), were split into “Convention” and “No Convention” based on whether they had reached a convention in Session 1. Within each of these groups the pairs were further divided into “Convention” and “No Convention” based on their performance in Session 2. This allowed us to evaluate which conditions were conducive to the formation, maintenance, or abandonment of a convention with a change in visual access.

#### 2.4.3. Phase II

##### Overall performance

Metrics for overall performance, and for the emergence and stability of conventions, were calculated for the Phase II pairs in the same manner as in Phase I Session 1, and compared to pairs in the instruction and opaque condition (I-O) of Phase I. We tested whether the proportion of pairs with conventions differed between Phases I and II using a Fisher’s exact test. A two-tailed t-test was used to determine whether the time taken for a convention to emerge and the stability of the conventions differed between I-O pairs from Phase I and Phase II pairs.

##### Transmission of conventions

For the pairs that had a convention emerge in Phase II, Spearman rank correlations were calculated between (i) the experienced subject’s color hierarchy at the end of Phase I, (ii) the color hierarchy they recalled at the beginning of Phase II, (iii) the color hierarchy developed by the experienced subject at the end of Phase II, and (iv) the color hierarchy developed by the naïve subject at the end of Phase II. These correlations were designed to test the extent to which conventions were transmitted across the phases and within Phase II. For each correlation of interest, the values were averaged across all the pairs to test for significance at the group level.

## 3. Results

### 3.1. Phase I Session 1

#### Overall performance

Conventions – consisting of a consistent color hierarchy used by both subjects of a given pair – emerged in all experimental conditions. The number of pairs that established conventions was 6 out of 10 in I-O, 3 out of 10 in I-T, 6 out of 10 in NI-O, and 4 out of 10 in NI-T. Thus, the opaque conditions contained more pairs with conventions (12/20) than the transparent conditions (7/20); however, a Fisher’s exact test showed that the difference among conditions was not significant (two-tailed, p = 0.538). Mean reaction time for pairs with a convention (mean ± SD = 1.71 ± 0.24 s, n = 9) did not differ significantly from those without a convention (mean ± SD = 1.34 ± 0.50 s, n = 11; t(18) = 2.03, p = 0.057); however, overall accuracy (percentage correct) in the task was significantly higher in pairs that used a convention (mean ± SD = 89.92 ± 8.81, n = 19) than in those that did not (mean ± SD = 76.43 ± 23.74, n = 21; t(25.9) = 2.36, p = 0.026).

#### Color hierarchies

We observed a variety of different color hierarchies across the 19 pairs that had developed a convention. Although some colors seemed to be preferred over others at the population level – for example, colors from the blue/purple end of the spectrum as well as pink were preferred over yellow, orange, and brown – we found weak agreement of color order across the pairs with conventions (Kendall’s W = 0.054, p = 0.056; see Supplementary Figure S2).

#### Emergence of conventions

The overall regression was statistically significant (R^2^ = 0.33, F(2, 16) = 5.38, p = 0.016), suggesting that the time taken for a convention to emerge varied between conditions. Visual access did not significantly predict time taken to reach a convention (β = −23.58, p = 0.392); however, instructions did (β = −75.26, p = 0.013). Of the pairs that reached a convention across both types of visual access (opaque and transparent), those with instructions (pairs that were explicitly told that choosing the same color would yield a reward) took longer (mean ± SD = 151.44 ± 65.28, n = 9) to reach a convention than those with no instructions (mean ± SD = 69.90 ± 45.73, n = 10; t(17) = 3.18, p = 0.005, Fig 2D).

#### Stability of conventions

Stability was only calculated for four of the six pairs with a convention in I-O since in two of the pairs the convention emerged late in the session, leaving fewer than 100 trials to analyze. For all other conditions, all pairs with conventions were used in the calculation of stability. The overall regression was statistically significant (R^2^ = 0.62, F(2, 14) = 13.87, p < 0.001), suggesting that the stability of conventions varied between conditions. Instructions did not significantly predict the stability of a convention (β = −0.005, p = 0.795), but visual access did (β = −0.088, p < 0.001). Of the pairs that had a convention across both types of instruction conditions, those with no visual access (opaque, mean ± SD = 0.993 ± 0.003, n = 8) had more stable conventions than those with visual access (transparent, mean ± SD = 0.969 ± 0.015, n = 9; t(15) = 4.47, p < 0.001, Fig 2E).

#### Self-reported strategies

The overall regression was statistically significant for minimum strategy used by the pair (R^2^ = 0.13, F(2, 36) = 3.88, p = 0.030), suggesting that condition may have had an effect on the minimum level of strategic behavior used by pairs. Visual access significantly predicted the minimum strategy used (β = 0.42, p = 0.01); however, instructions did not (β = 0.08, p = 0.630). Pairs in the transparent conditions had a higher minimum strategy (mean ± SD = 0.90 ± 0.45, n = 20); than those in the opaque conditions (mean ± SD = 0.47 ± 0.51, n = 19; t(35.73) = 2.76, p = 0.009). The overall regression was not statistically significant for maximum strategy used (R^2^ = −0.02, F(2, 36) = 0.64, p = 0.534), suggesting that condition did not affect the maximum level of strategic behavior elicited.

The regression model from the “Emergence of conventions” section was rerun for the pairs with both conventions and strategy data available (18 pairs, as opposed to 19 pairs with conventions previously), yielding similar significant results as above (R^2^ = 0.31, F(2, 15) = 4.90, p = 0.023). Adding minimum strategy as an independent variable did not improve the model (R^2^ = 0.28, F(3, 14) = 3.18, p = 0.057) and neither did adding maximum strategy as an independent variable (R^2^ = 0.28, F(3, 14) = 3.18, p = 0.057).

### 3.2. Phase I Session 2

#### Maintenance of conventions across conditions

All 10 pairs that had developed a convention in the opaque condition maintained a convention in their subsequent transparent session as well, while 4 out of 10 of the pairs that did not have a convention in the opaque condition developed one in the transparent. Meanwhile, a third of the pairs (3 out of 9) that had developed a convention in the transparent condition did not maintain one when they transitioned to their opaque session, and of the remaining 11 pairs that did not develop a convention during their transparent session, 7 developed one in the opaque session (Fig 2F).

To address the unexpected result of conventions being lost upon transitioning from a transparent to an opaque session, we hypothesized that instead of these pairs having established a convention in the form of an arbitrary *shared* hierarchy of colors, one subject was using a color hierarchy in the transparent session while the other was simply copying without being explicitly aware of the hierarchy. Once the opaque partition was introduced and copying was no longer possible, what appeared to be a shared color hierarchy disappeared since only one of the subjects had internalized it (see example pair in Supplementary Figure S3, showing the complete collapse of the hierarchy apparent in Subject 2’s Session 1 Elo-scores once visual access to the partner is removed for an opaque Session 2). This allowed us to categorize the 9 pairs with apparent color hierarchies in the transparent conditions of Phase I Session 1 (I-T and NI-T) into those with a “true” color-hierarchy convention (where both subjects had internalized the same hierarchy; 6 pairs) and those without (where this appeared to be the case for only one of the subjects; 3 pairs). We in turn found that reaction time per trial in the transparent first session for those that later maintained their convention was significantly faster than for those that lost it (maintained: mean ± SD = 1.69s ± 0.20s, n = 6; lost: mean ± SD = 2.14s ± 0.26s, n = 3); t(7) = 2.91, p = .023).

For the 6 pairs that continued to use a color-hierarchy convention, Spearman rank correlations revealed high levels of consistency between each subject’s color hierarchies at the ends of Sessions 1 and 2 (transparent and opaque, respectively), suggesting that they had indeed solved the initial transparent session via a *shared* color hierarchy (median correlation coefficient: 0.80). Importantly, the effect of condition on the time taken for the emergence of conventions held even when the three pairs without a true color hierarchy convention were excluded from the analysis: the model yielded similar significant results as the full dataset of 19 pairs (R^2^ = 0.46, F(2, 13) = 7.54, p = 0.007; n = 16 pairs). Instructions remained a significant predictor of time taken to reach a convention (β = −97.36, p = 0.003), with pairs with instructions taking longer, i.e. more trials to reach a convention, (mean ± SD = 165.75 ± 52.58, n = 8) than those with no instructions (mean ± SD = 66.00 ± 47.13, n = 8; t(13.84) = 4.00, p = 0.001).

### 3.3. Phase II

#### Overall performance

Of the 21 tested pairs in Phase II, 13 developed a convention. This proportion (13/21) did not significantly differ from that observed in Phase I Session 1 pairs (I-O condition: 6/10, p = 1.000). However, Phase II pairs reached a convention significantly faster than pairs in Phase I (I-O) (Phase II: mean ± SD = 121.46 ± 38.83 trials; Phase I: mean ± SD = 174.5 ± 59.19 trials; t(17) 2.37, p = 0.030; see Fig 3A). There was no significant difference between the stability of conventions in Phase II (mean ± SD = 0.993 ± 0.004, n = 13) and pairs in Phase I Session 1 I-O that had a convention (mean ± SD = 0.995 ± 0.002, n = 4; t(15) = 0.65, p=0.526, Fig 3B).

**Figure 3.**
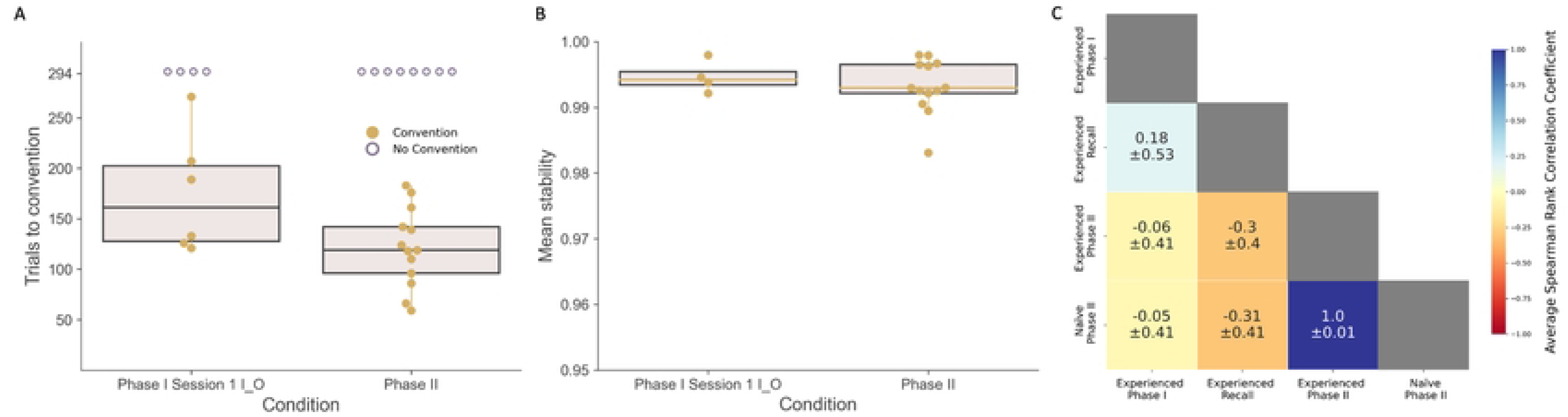
Phase II Results: Transmission of Conventions. (A) Trials taken to reach a convention in the pairs that had a convention (solid orange dots) in Phase I (Session 1 I-O condition; 6/10) compared with those in Phase II (13/21). Hollow purple dots indicate pairs that did not converge on a convention and were not included in the calculation of the means for each condition and (B) mean stability of the conventions that emerged in Phase I (Session 1 I-O condition) compared with mean stability of conventions in Phase II. For A and B, upper and lower bounds of box represent upper and lower quartile, respectively, center of the box represents the median and the whiskers extent to maximum and minimum (outliers are calculated ± 1.5 * inter quartile range past the high and low quartiles). (C) Mean correlations across pairs in Phase II that had a convention between (i) Experienced – Phase I: the experienced subject’s color hierarchy at the end of Phase I, (ii) Experienced – Recall: the color hierarchy they recalled at the beginning of Phase II, (iii) Experienced – Phase II: the color hierarchy developed by the experienced subject at the end of Phase II, and (iv) Naïve - Phase II: the color hierarchy developed by the naïve subject at the end of Phase II.

#### Transmission of conventions

Figure 3C summarizes all correlations among experienced subjects’ Phase I color hierarchies, their recalled hierarchies, their Phase II hierarchies, and the hierarchy developed by the naïve subject in Phase II. Only 2 out of the 21 experienced subjects had significant recall correlations (meaning they were able to remember their previous hierarchies from the end of Phase I; ρ(7) = 0.89, p = 0.007 and ρ(7) = 0.86, p = 0.013). Neither these two subjects, nor the set of experienced subjects as a whole, transmitted their Phase I hierarchies to the naïve subjects: we found no significant mean Spearman rank correlations between the experienced subjects’ hierarchies at the end of Phase I (or their recalled hierarchies) and their naïve partners’ hierarchies at the end of Phase II. Only one pair had a significant correlation between the experienced subject’s recalled hierarchy and the transmitted hierarchy, but, interestingly, this was a negative correlation (ρ(7) = - 0.89, p = 0.007).

## 4. Discussion

The use of conventions – arbitrary, self-sustaining practices that emerge in a population and facilitate efficient solving of a specific coordination problem – are a ubiquitous aspect of human behavior and communication [7,8]. Nevertheless, many open questions remain surrounding the process through which conventions are established and maintained, the level of cognitive sophistication required, as well as the group-level processes that facilitate their emergence. Our study demonstrates the emergence of simple conventions in experimental human pairs for solving a novel coordination problem presented across a number of conditions, even in the absence of communication. Specifically, we examined how participants’ knowledge of the structure of the problem, their access to their partner’s actions, and their prior experience with solving the task affected whether and how quickly the conventional solution was found by a given pair. We further analyzed the stability of the conventions formed as a function of these conditions as well as the transmission of conventions from knowledgeable to naïve individuals (simulating how established conventions may be conserved under demographic turnover in a hypothetical population). Strategies reported by the subjects at the end of their session were used to both quantitatively and qualitatively assess the cognitive processes that may have been recruited for the successful completion of the task, as well as other conventions that may have emerged (see below).

Crucially, all three key features of conventions (efficient, arbitrary, self-sustaining) were evident in our data. First, we were able to confirm that pairs with a convention outperformed those without, achieving higher overall scores. Second, we observed a number of different color hierarchies, with no statistically significant agreement across pairs in the ranking of the colors (although some colors did appear to be highly ranked by more pairs than others, possibly due to salient features such as brightness), demonstrating the arbitrariness of the solutions that emerged. Third, although some pairs that developed a convention in their first session lost it following a change in visual access to the partner’s actions (second session), the majority (16 out of 19) maintained theirs even across this shift.

Our Phase I Session 1 results also indicate, contrary to our expectations, that having more information about the structure of the problem does not necessarily increase the speed with which conventions are first established. In fact, we observed the opposite of such an effect: in the two conditions where our subjects had explicit instructions that choosing the same color would yield a reward (I-O and I-T), i.e., they knew what counted as a case of successful coordination from the start, conventions took longer to emerge. We hypothesize that having more information about the task structure could have resulted in the subjects paying more attention to their partner’s actions (when visible), as well as potentially building mental models of their partner’s strategy (whether or not the partner’s actions were visible). On the other hand, a simpler reward-contingency-based approach, in which subjects repeatedly sample the environment and build up expectations about the relative payoff likelihoods of the different colors, without forming any beliefs about their partner, could have yielded faster convergence on a conventional solution in the conditions with no instructions (NI-O and NI-T). Indeed, such a strategy likely explains baboon subjects’ convergence on a hierarchy of stimuli in Formaux and colleagues’ [9] study, most saliently in the condition where monkeys had no visual access to their partner’s choices at any point during learning (experiment 2.2). Our hypothesis to explain our results is in line with findings in the game theory literature where reinforcement learning leads to faster convergence to a solution compared to more complex strategies that attempt to predict the partner’s behavior (e.g., [33–35]). However, our analysis of the levels of strategic reasoning employed by subjects – as reflected in their self-reported strategies – did not support this hypothesis, in that we found that condition (I vs NI) did not significantly predict subjects’ level of strategic reasoning. Nonetheless, we recommend caution in the interpretation of these results since subjects’ strategies were obtained from a free response questionnaire where different subjects responded with varying levels of detail. To better tease apart underlying mechanisms, a more specific post-session questionnaire would be required, carefully crafted so as not to bias subjects into reporting, for example, more complex strategies (via retrospection) than the one that they actually employed during the task.

In contrast with our speed-of-emergence comparisons, the *stability* of conventions did not vary as a function of the information provided to subjects at the beginning of the experiment. It did, however, vary with visual access: conventions in opaque conditions (I-O and NI-O) were more stable than those in transparent conditions (I-T and NI-T). This was as expected: in opaque conditions a color-hierarchy convention was the only way to solve the task, whereas in transparent conditions subjects could instead also use social information trial-by-trial (i.e., check the partner’s screen for their choice and copy it), likely leading to the less stable color hierarchies we observed. In addition, visual access also appeared to influence the level of strategic reasoning elicited in the subjects: the minimum strategic level employed by pairs was higher in the transparent conditions than in the opaque conditions. This was surprising given the availability of the copying solution in the transparent conditions but may suggest that visual access to a partner’s actions prompted the engagement of higher-level processing of social information. If accompanied by higher responsivity to the partner’s actions (including to any choices made in error, i.e., not matching a currently established color hierarchy), then such higher-level reasoning about the partner may, in turn, have been another factor responsible for the increased changeability (lower stability) of color hierarchies in the transparent conditions.

Indeed, the availability of a copying strategy under the transparent conditions represents an important caveat. While a shared hierarchy of colors is one possible convention that enables fast and consistent solving of the task, division of labor or turn-taking are also kinds of conventions that can serve as solutions to coordination problems (e.g., [27]). For example, in our task, an alternative convention in the transparent conditions would see one of the subjects emerging as the leader or “proposer”, i.e., the one who consistently chooses one of the two options first, and the other becoming the follower or “responder”, who then chooses the same color as the proposer (this was in fact an integral aspect of the design in Formaux and colleagues’ [9] study with baboons, although there the roles were randomly assigned for each session). This strategy is efficient since the consistency in roles removes the uncertainty about who should choose first on each trial and thus neither partner is delayed in attending to their half of the task (choose color for the proposer, and copy partner for the responder). As such, this alternative strategy also carries the hallmarks of a convention: not only is it efficient, but it is also arbitrary in that either player could assume either role, and it is self-sustaining in that an attempt by one subject to reverse the roles would interfere with the efficient solving of the task (at least temporarily). However, it is unclear how such a convention would scale up to the population level: coordination would emerge quickly whenever a proposer and a responder happen to meet, but two proposers or two responders would experience difficulty. A color-hierarchy-based convention, as long as it is shared by the population, would ensure efficient solving between any two randomly chosen individuals from the population (the nationally mandated driving conventions referenced in the Introduction represent a clear example).

We were not able to determine from the trial-by-trial response data which pairs adopted a proposer-responder convention (with or without also establishing a shared or unshared, i.e., proposer-only, color hierarchy): in theory, reaction times could reveal which player chose first, but in practice we observed that proposers tended to move their cursor over the color they were about to choose, waited for the responder to also move their cursor over the same color, and then both subjects clicked simultaneously, leading to near-identical reaction times. Nonetheless, two other lines of evidence do suggest the emergence of such a convention in some pairs. First, strategies reported by subjects in transparent conditions (across both I and NI) included explicit mentions of a given subject “following” the choices of their partner consistently. Second, three pairs exhibited a curious pattern where they appeared to have generated a color hierarchy convention in their transparent Session 1 (where a convention was not essential for solving the task) but immediately lost it in the following opaque Session 2 (where it was necessary). It is likely that instead of these pairs having established a convention in the form of an arbitrary *shared* hierarchy of colors, one subject was using a color hierarchy in the transparent session while the other was simply copying without being explicitly aware of the hierarchy. Interestingly, we found that pairs with “true” color conventions were faster at the task at the trial level than these pairs. This suggests that within the framework of our task, the color-hierarchy convention was not only more robust to changes in context (presence vs absence of visual access) than the proposer-responder convention, but also overall more efficient in a context where both types of convention were available to subjects.

These varied outcomes in how pairs approached our task allow us to speculate about the role of specific cognitive mechanisms that support the different kinds of conventions available to subjects. We argue that the establishment of a shared color hierarchy between players required, at minimum, the ability to track reward contingencies obtained through environmental sampling over time (where these reward contingencies are, unbeknownst to one player, being modified in real time by the other player). In fact, this ability may be a necessary and sufficient condition enabling the gradual emergence of a shared color hierarchy, extending even to cases where subjects are not aware of the existence or role of a partner. In contrast, a strategy involving copying of the partner may require additional social cognitive skills: at minimum it relies on the capacity for social information use and learning to map the appropriate response to the partner’s observed choice, but can also be supported by more complex forms of ‘cognitive imitation’ [36] where the subject copies a cognitive rule that it infers the partner is using. Copying skills are prominent in humans [37,38], and while they are likely also present among non-human primates, monkey and ape subjects typically require extensive training before they are able to use the choice behavior of another individual as the basis for their own choice [36,39]. While baboon pairs in Formaux and colleagues’ [9] study could not spontaneously develop a proposer-responder convention given that the experimental design automatically assigned one baboon as the proposer and the other as the responder on every trial, they were observed to watch their partner before responding only when the two colors had similar Elo-scores (and hence the trial was “difficult”). In contrast, over a third of our pairs in transparent conditions developed a proposer-responder convention reliant on copying.

Moreover, our human pairs also leveraged other cognitive abilities: for example, despite our instructions specifying that subjects were not allowed to communicate with each other, behaviors such as hovering over the chosen stimulus until the other player’s cursor arrived at the same color arguably circumvented this constraint. In addition, subjects’ self-reported strategies also revealed a tendency to attend at least to the choices of the other player and, in some cases, to take this further by speculating about the other player’s strategy and future behavior based on their prior choices. Such modelling of the other player’s mind would appear to recruit advanced socio-cognitive skills (such as theory of mind) potentially unique to humans. Interestingly, we suspect that one of the reasons for the delayed emergence of color hierarchy conventions in our Instruction conditions was that knowing more about *how* the partner’s choice influenced trial outcomes prompted the increased engagement of high-level social cognition. Specifically, this engagement could have led to oscillations in players’ choices as each player attempted to “catch up” with their partner (e.g., on an “incorrect” trial, both players may infer that the other player selected the other color leading both to adjust their own choice, with the same incorrect outcome on the next trial and hence both shifting their choice again, and so on). Hence, somewhat counterintuitively and potentially as an artefact of our experimental design prohibiting overt communication, we found that simpler cognitive strategies likely enhanced the speed with which conventions emerged.

Exploring the minimum cognitive underpinnings required for different types of conventions further allows us to hypothesize where and under what circumstances we may expect to find conventions used by non-human animals. Taking cooperative chick rearing among birds where parents take turns at the nest as an example, we can ask whether we could in theory expect given pairs to settle on a specific daily ‘handover’ time for chick duties. These could be considered conventions if the timing is indeed at least somewhat arbitrary (i.e., switching over at 2pm rather than 4pm has no impact on chick rearing success), efficient (i.e., each parent can maximize its own foraging time without risking the chick being left on its own), and self-sustaining (i.e., once a specific handover time is ‘agreed’, any deviation from it will disadvantage at least one of the parents as well as the chicks). How might such a convention emerge? If we consider arriving back to the nest just as the other is preparing to leave a ‘success’, then any arrivals that miss this window could prompt an adjustment of the return time until there is a match; similarly, a parent who decides to leave before the arrival of the other may leave later in future (hunger permitting) until a window of overlap is found. This process would not require any advanced social cognition, either in terms of predicting the partner’s likely future behavior or in modelling the ‘strategy’ that the partner may be using. Although this example is highly speculative and would require longitudinal data on the development of handover times in newly mated pairs to verify, it leads us to hypothesize that conventions among non-humans in nature may emerge more frequently than currently realized.

Phase II of our study was designed to simulate the transmission of conventions from knowledgeable individuals to those ‘naïve’ to a population’s or their partner’s existing conventions. In multigenerational populations and in fission-fusion societies, for example, such naïve individuals may arrive via birth, immigration, or a reshuffling of subgroup membership. Our results showed that conventions emerged faster when one of the subjects was experienced (i.e., had learnt to solve the task via a memorized color hierarchy developed in their previous pair) but that this did not happen through the faithful transmission of the *same* solution that had previously emerged. Specifically, it appeared that knowledgeable individuals were able to transmit know-how (i.e., a hierarchy of colors being necessary) but not know-what (i.e., the specific ranking of colors in such a hierarchy). Such effects may be particularly prominent in cases where interactions are purely dyadic and repeated interactions with new partners are rare or non-existent (such as in the case of the chick-rearing example discussed above). Nonetheless, since we also found that our knowledgeable subjects failed to accurately recall at the start of Phase II their color hierarchies established during Phase I, it remains possible that they did not have sufficiently ‘engrained’ know-what information to transmit it accurately. Future work should examine the role of population turnover rate (affecting the proportions of naïve and knowledgeable individuals available in the population), interaction frequency (affecting the period over which the solution has to be maintained in memory without reinforcement), interaction type (whether interactions occur primarily with a fixed partner or more freely across the population), individual differences in flexibility to adjust existing preferences (affecting how likely, for example, a stubborn minority would be to accommodate population-level shifts), and the cost of failed interactions on the stability of both the form and specific content of conventional solutions.

Conventional behavior as an efficient solution to coordination problems forms the basis of many of our interactions as humans. Understanding how conventions emerge, under what conditions they are maintained stably, and what might cause them to change or disappear, can have significant implications on the functioning of our large, complex, and interconnected societies. In future, results like ours could help inform how we can best present information to individuals in a way that allows for rapid convergence to conventional solutions to coordination problems, increasing the efficiency of our interactions. Future studies should also address how these conventions are transmitted between knowledgeable and naïve individuals in a population, how other factors like group size, problem context, cost of failure to coordinate, or frequency of interactions shape the formation and maintenance of conventions, and how pairwise interactions could best be structured to allow quick convergence to conventions at the population level.

## 5. Acknowledgements

We thank Kristin Andrews, Nicolas Claidiere, and Robert Hawkins for helpful comments on an earlier draft of this manuscript. We are also grateful to Enana Jacob for help with coding subjects’ self-reported strategies.

## 7. Supporting information

**S1 Fig.** Two complete example pairs from Phase I Session 1 I-O.

Left (A-D) for a pair with a convention and right (E-H) for a pair without a convention

Evolution of Elo-ratings for Subject 1 (A) and Subject 2 (B) of an example pair that developed a convention. (C) Mean combined accuracy as a function of trial number, illustrating the emergence of a convention for an example pair. Combined accuracy is a metric that takes into account (i) whether the two subjects chose the same colour *and* (ii) whether both subjects’ choices matched their respective colour hierarchies developed up to that point. Mean combined accuracy is calculated over rolling windows of 21 trials. Note that the plot starts at trial 21 and each data point represents the mean of the previous 21 trials. Horizontal green line indicates threshold above which pairs had to maintain performance for at least 21 consecutive overlapping windows in order to pass criterion; vertical yellow line indicates the trial at which this pair was judged to have passed criterion, i.e., where a shared colour hierarchy emerged and was maintained between the two subjects.

(D) Elo-ratings of the two subjects plotted together across the session for every set of 42 trials (this included all possible colour combinations being presented exactly twice). Increasing size of the circles correspond to progression through 7 such sets of 42 trials (yielding a total of 294). The final Elo-rating values at the end 294 trials are plotted with a black outline and the black line represents the regression through these points.

E-H show the same plots for a pair without a convention.

**S2 Fig.** Mean ranking of the seven colours across subjects in pairs that had developed a convention in Phase I Session 1. Lower rank number implies the colour appeared at the top of more hierarchies. Error bars show SEM (standard error of the mean).

**S3 Fig.** Example pair’s evolution of Elo-ratings across both sessions of Phase I. This example pair completed the first session in NI-T condition followed by an opaque session. Analysis of Session 1 (transparent) shows a convention emerging in the pair but looking at Session 2 (opaque) shows us that Subject 1 (left) may have been using a color hierarchy to solve this task while Subject 2 (right) was likely copying Subject 1 in Session 1. What appeared to be a convention the transparent session disappeared in the following opaque session.

**S1 Table.** Hypothetical and subject-reported examples of Level 0, 1, and 2 strategies.

